# Unravelling resilience mechanisms in forests: role of non-structural carbohydrates in responding to extreme weather events

**DOI:** 10.1101/2020.09.03.281105

**Authors:** Ettore D’Andrea, Andrea Scartazza, Alberto Battistelli, Alessio Collalti, Simona Proietti, Negar Rezaie, Giorgio Matteucci, Stefano Moscatello

**Affiliations:** CNR-ISAFOM, P.le Enrico Fermi 1 - Loc, Porto del Granatello, 80055 Portici NA, Italy; CNR-IRET, Via Moruzzi 1, 56124 Pisa, Italy; CNR-IRET, via Marconi 2, 05010 Porano (TR), Italy; CNR-ISAFOM, Via della Madonna Alta 128, 06128, Perugia (PG), Italy; DIBAF, University of Tuscia, 01100 Viterbo, Italy; CREA-IT, Via della Pascolare, 16, 00015, Monterotondo Scalo (Roma), Italy; CNR-IBE, via Madonna del Piano, 10 - 50019 Sesto Fiorentino (FI), Italy

**Keywords:** Carbon allocation, Carbon reserves, Drought, Fagus sylvatica L., Late frost, Mediterranean, Phenology, Resilience

## Abstract

- Extreme weather events are increasing in frequency and intensity due to global climate change. We hypothesized that these have a strong impact on the stem radial growth and the dynamic of non-structural carbohydrates (NSCs).
- In order to assess the effects on mature trees of a late frost occurred in spring 2016 and a drought event characterizing the summer 2017, we monitored the phenology, the radial growth and the dynamic of starch and soluble sugars in a Mediterranean beech forest.
- Growth was much more reduced by spring late frost than by summer drought, while NSCs dynamic was deeply involved in counteracting the negative effects of both events, supporting plant survival and buffering source-sink imbalances under such stressful conditions, resulting in a strong trade-off between growth and NSCs dynamic in trees.
- Overall, our results highlight the key role of NSCs on trees resilience to extreme weather events, confirming the relevant adaptability to stressful conditions. Such an insight is useful to assess how forests may respond to the potential impacts of climate change on ecosystem processes and to define how future management strategies can help adaptation of beech forests in the Mediterranean area.

## Introduction

Global climate change is causing an increment in the frequency of extreme weather events (Stocker *et al.*, 2014) that are recognized among the major drivers of current and future ecosystem dynamics (Frank *et al.*, 2015). The Mediterranean basin is one of the two main hot-spots of climate change (Giorgi, 2006; Noce *et al.*, 2017), showing increases in the inter annual variability and of extreme environmental conditions (Flaounas *et al.*, 2013). In this region, the increasing risk of late frost events represents one of the major threats associated with the future global change (Zohner *et al.*, 2020). Indeed, increasing spring temperatures has been observed stimulating earlier leaf unfolding (Gordo & Sanz, 2010; Allevato *et al.*, 2019), thus potentially exposing young leaves and shoots to spring frost damage (Augspurger, 2013), especially at high elevation (Vitasse *et al.*, 2018). Depending on species, temperatures below −4°C can destroy the “fresh” leaves and shoots reducing - up to even blocking - the photosynthetic capacity of trees for several weeks. In this case, the resource requirements for new leaves formation, and tree life maintenance, must necessarily rely on the remobilization of carbon (C) reserves (Dittmar *et al.*, 2006; D’Andrea *et al.*, 2019). Moreover, severity, duration, and frequency of drought events have all been increasing in the last decades (Spinoni *et al.*, 2015). European beech (*Fagus sylvatica* L.), one of the most diffused native tree species in Europe, is known to be drought sensitive (Bolte *et al.*, 2016). Hence, drought events can negatively affect physiological performance (Rezaie *et al.*, 2018), carbon allocation (D’Andrea *et al.*, 2020a), reproductive capacity (Nussbaumer *et al.*, 2020), as well as the growth and competitive strength of the species (Peuke *et al.*, 2002) which may all impact its future distribution (Noce *et al.*, 2017).

Growth and non-structural carbohydrates (NSCs; i.e. sucrose, fructose, glucose and starch) dynamic are among the most strongly affected ecosystem processes by spring frost and summer drought (Li *et al.*, 2018). An increasing body of evidence has shown that NSCs dynamic is not a pure passive deposit and removal of C compounds, but it represents a key process actively controlled by plants to finely regulate C source-sink balance and to buffer the difference between C supply and demand at different timescales (Scartazza *et al.*, 2001; Sala *et al.*, 2012; Carbone *et al.*, 2013; Fatichi *et al.*, 2014; Moscatello *et al.*, 2017; Collalti *et al.*, 2020). Therefore, NSCs could play a crucial role in counteracting the negative effects of extreme weather events on beech forests, contributing to their resilience and survival (Scartazza *et al.*, 2013; D’Andrea *et al.*, 2019). Unfortunately, despite the recognized importance of NSCs for plant productivity and resilience, little is known regarding their seasonal regulation and trade-off with growth and reproduction in forest trees (Merganičová *et al.*, 2019; Tixier *et al.*, 2020).

In this work, we studied the effects of spring late frost and summer drought in a long-term research plots established on a Mediterranean beech forest of Central-South Italy (Collelongo, Abruzzi Region, Italy). The site is located in the large area in the Central-South Italy where in spring 2016, due to unusually warm preceding weeks, leaf unfolding occurred up to 15-20 days earlier than the normal average, followed by frost, that caused the complete loss of the newly grown canopy. Moreover, in 2017, a strong summer drought, due to a combination of drastic reduction of precipitation associated with high air temperature in late July and August, interested a huge area of the Mediterranean basin (Bascietto *et al.*, 2018; Nolè *et al.*, 2018; Allevato *et al.*, 2019; Rita *et al.*, 2019), including the Collelongo site. Notwithstanding these events were monitored through remote sensing techniques, *in situ* evaluations of their effects on ecosystem functionality are limited. Phenology, growth and NSCs dynamic in the Collelongo beech stand were investigated during 2016 (i.e. the year of the late frost event) and 2017 (i.e. the year of the summer drought event) and compared to the historical inter-annual data collected earlier at the site.

The objectives of the study were to: *i*) quantify the magnitude of the effects of such extreme events on ecosystem functioning; *ii*) verify the role of NSCs in mediating source-sink balance following the strong alteration of C activity; *iii*) evaluate the interplay and trade-off between carbon allocation to canopy, stem and C reserves. The aim was to predict how these responses and regulation processes could contribute to the resilience of beech to extreme weather events associated with future global change.

## Material and methods

### Study site

The study was carried out during the years 2016 and 2017 in an even-aged, pure beech forest (*Fagus sylvatica* L.) located at Selva Piana stand (41°50’58” N, 13°35’17” E, 1560 m elevation) in the Central Apennine (Collelongo, Abruzzi Region, Italy). The experimental area is included in the LTER network (Long Term Ecological Research) and more specific information about the site were already reported in previous works (Guidolotti *et al.*, 2013; Collalti *et al.*, 2016; Rezaie *et al.*, 2018; Reyer *et al.*, 2020) in which the soil, forest structure, and climate characteristics are described.

### Climate and Phenology

The temperature and precipitation for the period 1989-2015, available on the Fluxnet2015 release, were used to characterize the, on average, climate conditions of the site. For the data gaps occurred during the experimental trial (2016-2017), we used the ERA5 database produced by the European Centre for Medium-Range Weather Forecasts (ECMWF) (https://www.ecmwf.int/en/forecasts/datasets/archive-datasets/reanalysis-datasets/era5, data accessed: [12/04/2018]), according to the Fluxnet2015 release formulations (Pastorello *et al.*, 2020). To evaluate peculiarities of the climatic conditions in 2016 and 2017 we calculated monthly differences with respect to the average values of precipitation and temperature observed in the site in the historical time series 1989-2015.

Leaf phenology was monitored using the MODIS Leaf Area Index product (LAI, MOD15A2H product, https://modis.gsfc.nasa.gov/) with 8-day temporal resolution and 500-meter spatial resolution (Myneni *et al.*, 2015). Critical dates, representing approximately linear transitions from one phenological phase to another, were identified and defined according to Zhang *et al.* (2003) as: (1) *green-up*, photosynthetic activity onset; (2) *maximum LAI*, supposed to be the leaf maturity phase; (3) *senescence*, sharp decrease of photosynthetic activity and green leaf area; (4) *winter dormancy.* In 2016, the leafless period after the late frost was identified from the day of the extreme event and the second green up.

### Selection, measurements and sampling of trees

Five trees were selected according to their similarity with site tree ring chronology, the trees had diameters at breast height (DBH) ranging from 49 to 53 cm (for more details see D’Andrea *et al.* 2020b). Trees were monitored from April 2016 to November 2017. Intra-annual radial growth of each selected tree was measured using permanent girth bands with 0.1 mm accuracy (D1 Permanent Tree Girth, UMS, Germany). Furthermore, stem diameter was recorded at the moment of each sampling of xylem for biochemical analyses (20 sampling dates from April 2016 to November 2017).

From each tree, micro-cores (2 mm diameter, 15 mm long) of wood were collected after bark removal, using the Trephor tool (Rossi *et al.*, 2005). All samples for biochemical analyses were immediately placed in dry ice for transport to the laboratory, then stored at −20 °C and, finally, stabilized through lyophilisation processes until NSCs analysis.

Daily radial increment (R_i_, μm day^−1^), was calculated as follow:

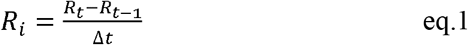

where *R* is the radius of each *i* tree (μm), *t* is the date of sampling, and *Δt* is the time interval between the two sampling dates expressed in days.

In November 2017, at the end of the experimental trial, increment cores were collected at breast height from each tree, using an increment borer. Tree ring width series were converted into tree basal area increment (BAI, cm^2^ year^−1^), according to the following standard formula:

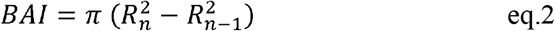

with *n* being the year of tree-ring formation.

### Starch and soluble sugar concentrations analysis

The freeze-dried xylem samples were milled to a fine powder and used for all analytical tests. For analysis of glucose, fructose, sucrose and starch, 10 mg of dry xylem powder were extracted in 1 ml of 80% ethanol/water at 80 °C for 45 minutes. After centrifugation at 16,000 x g for 5 minutes, soluble sugars were recovered in the supernatant while the pellet was resuspended in 1 ml of 40 mM acetate buffer (pH 4.5), then re-centrifuged 16,000 x g for 5 minutes. This procedure was repeated 4-times. The final pellet was autoclaved for 45 minutes at 120 °C in the same wash buffer. Enzymatic starch hydrolysis and the following glucose spectrophotometric assay were done as described by Moscatello *et al.* (2017). The supernatant solution containing soluble sugars was filtered on 0.2 μm nylon filters (GE-Whatman, Maidstone, UK), then analyzed by high-performance anion exchange chromatography with pulsed amperometric detection (HPAEC-PAD) (Thermo Scientific™ Dionex™ ICS-5000, Sunnyvale, CA U.S.A.)(Proietti *et al.*, 2017).

### Modelling of Intra-annual dynamics of non-structural carbohydrates

To evaluate the effects of the spring late frost (2016) and the heat wave and drought stress (2017) on the intra-annual NSCs dynamic, a representative benchmark of the typical intra-annual carbohydrates dynamic of the study site was needed. With this aim, a dataset on NSCs dynamic derived from other experimental trials at the site was used (Supporting Information Table S1). Dataset was composed of data of different years (i.e.: 2001, 2002, 2013, 2014, 2015, and 2018). This dataset included 39 observations of starch dynamic and 28 observations for both soluble sugars (glucose, fructose and sucrose) and total NSCs dynamic. Observations for soluble sugars were lower, because of the methodological sampling procedure used in 2015. During that year, woody samples were collected for xylogenesis analysis and maintained in ethanol-formalin acetic acid solution (FAA). Unfortunately, this methodology caused the loss of soluble sugars, while the starch integrity was preserved, as verified by means of specific analytical tests on woody tissues.

Different models based on data of starch, soluble sugars and total NSCs were used looking for possible patterns within the years and tested through the Akaike Information Criterion (AIC) (Akaike, 1974; Aho *et al.*, 2014) to select the simplest model able in reproducing the *in situ* observed pattern. The AIC quantifies the trade-off between parsimony and goodness-of-fit in a simple and transparent manner, estimating the relative amount of information lost by a given model. Hence, the model showing the lowest AIC is considered the model with the smallest information loss and, potentially, the most representative one (Akaike, 1974). The four assumptions of linear model (homoscedasticity, normality of the error distribution, statistical independence of the errors and absence of influential points) were tested graphically (Fig. S1). Statistical analysis and figures were made using R 3.5.0 (R Development Core Team, 2018).

## Results

### Climate in the study period

Monthly variations of temperature and precipitation in the Selva Piana beech forest are reported in Figure 1a-b. In 2016 a severe late frost event occurred during the night between April 25 and 26, when the temperature at canopy level (~ 24 m) reached – 6 °C (Fig. 1a inset panel). The extreme frost event followed an early spring season characterized by a temperature that during the months of February and April was significantly higher (about 2°C) than the average value of the site for the period 1989-2015 (Fig. 1a). In 2017, from May to August, the temperature was significantly higher than the average value of the site, with an increase of ~3 °C (Fig 1a). Furthermore, from May to October 2017 a significant reduction of precipitation against long term average was observed (Fig. 1b), leading to an annual precipitation that was ~ 50% of the 1989-2015 average (Fig. 1b inset panel).

**Figure 1:**
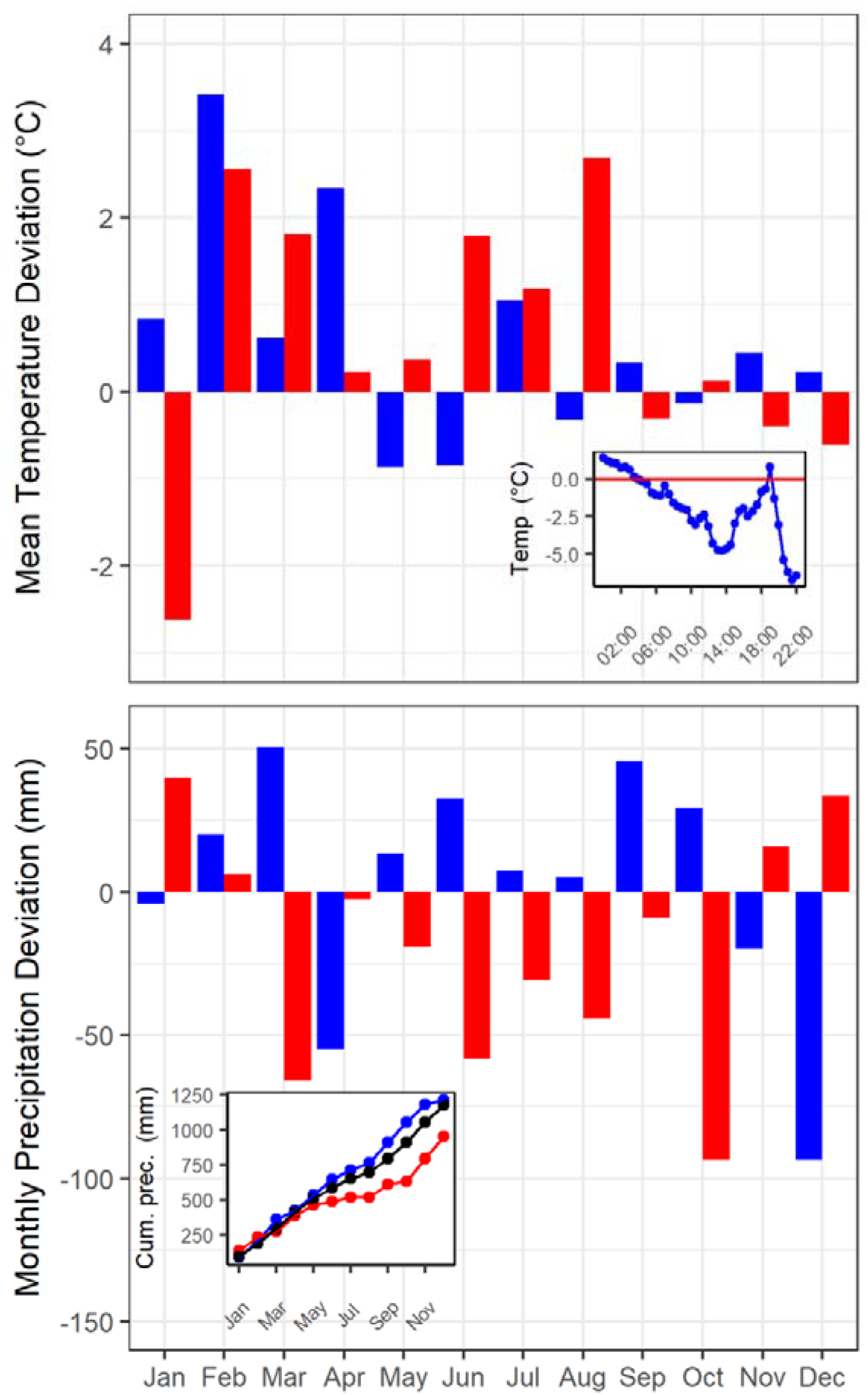
Deviations of monthly mean temperature and precipitation for 2016 (blue bars, panel A) and 2017 (red bars, panel B) calculated as the difference from the 2000-2015 average value at the site. Temperature of April 25^th^, 2016 measured at canopy level (24 m) is reported in inset graph of panel *a*, while the annual precipitation of the year 2016 (blue dots), 2017 (red dots) and the long-term average (black dots) is reported in the inset graph of panel *b*.

### Phenological parameters and radial growth

The seasonal LAI trend, used to define the phenological phases of the stand, is reported in Fig. 2a. The “first” green up in spring 2016 occurred between 20 and 30 days earlier than the average of the site (Fig. 2a), while the “second” (re)green up, after the complete canopy destruction due to the spring frost event, started around June 28, with a leafless period of more than 60 days. In 2016 the beginning of the senescence phase was anticipated of about one week compared to the average of the long-term series (Fig. 2b). Maximum LAI was lower in 2016 (LAI = 4.79 m^2^ m ^2^) than in 2017 (LAI = 5.37 m^2^ m^−2^), while the long-term average LAI of the site assessed with remote sensing was ~5 m^2^ m^−2^ (Fig. 2a). The average length of vegetative period assessed through remote sensing during the 2000-2015 period was approximately 140 days, a value confirmed in 2017, while it was 83 days in 2016.

**Figure 2:**
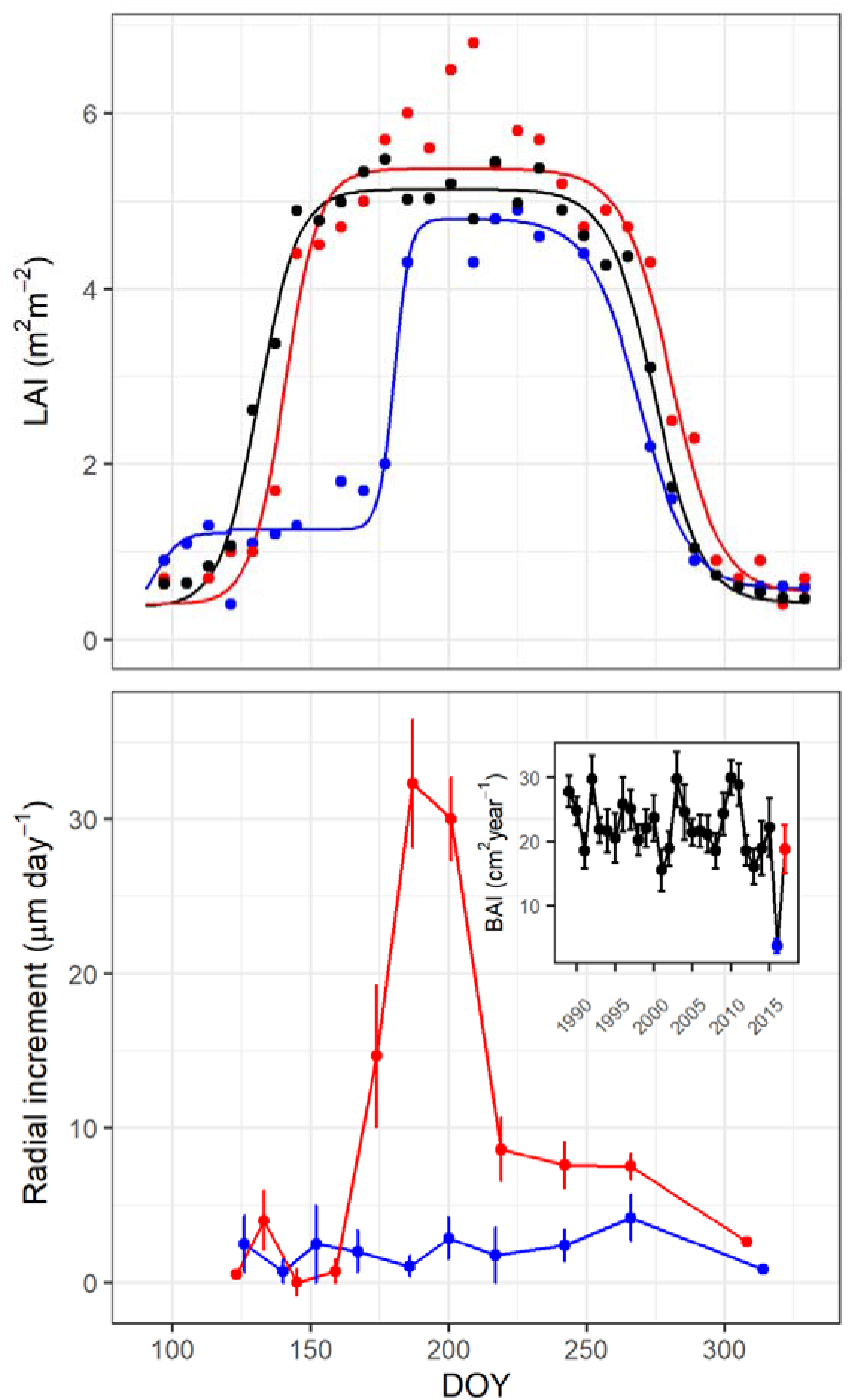
Seasonal dynamics of Leaf area index (LAI, m^2^ m^−2^, panel *a*) and daily stem radial increment (panel b) for the years 2016, 2017 and the 2000-2015 reference period. LAI was derived from Moderate Resolution Imaging Spectroradiometer (MODIS, see Materials and Methods), for 2016 (blue line), 2017 (red line) and for the 2000-2015 reference period (black line). Solid lines are the modelled LAI pattern, using two logistic functions for the increasing and decreasing phases. Dots are the raw MODIS-LAI values. In panel *b* the daily radial increment for 2016 (blue dots) and 2017 (red dots) are shown, while the inset graph reports the long-term series data of Basal Area chronology (BAI, cm^2^ year^−1^), where the last two dots represent the BAI value obtained in 2016 (blue dot) and 2017 (red dot), respectively.

The mean BAI in the 2000-2015 period was 22.64 ± 0.78 cm^2^ year^−1^, while it was 3.69 ± 1.14 cm^2^ year^−1^ and 18.75 ± 3.80 cm^2^ year^−1^ in 2016 and 2017, respectively (Fig. 2b inset panel). The late frost in spring 2016 reduced the stem radial growth of about 85% compared to the average of the period 1989-2015. The late frost strongly affected the seasonal dynamic of stem diameter growth during the year 2016, as shown by the lower and almost constant rate of stem growth compared to 2017, when after the green up the radial growth followed the usual pattern, reaching the highest increment (32.30 ± 4.14 μm day^−1^) in July (Fig. 2b).

### Intra-annual dynamic of NSCs

The values and the modelled intra-annual dynamics of NSCs (total sugars, starch and soluble sugars content) measured in the beech stem wood are reported in Fig. 3 (panels a, b and c). Dynamic of NSCs showed polynomial equation patterns at different grades, with R^2^ ranging from 0.64 to 0.93 (Table 1). Comparing the modelled NSCs intra-annual dynamics and stand phenology, an increase in total NSCs is observed from the bud break to the beginning of green-up phase, due to the increasing starch content notwithstanding the decrease of soluble sugars. During the period between the onset and the middle of the maximum vegetative season, total NSCs content decreased due to starch reduction, while the amount of soluble sugar remained unchanged. In the late summer, both starch and soluble sugars increased until the end of the vegetative season, determining an increase of total sugars content (Fig. 3a). At the beginning of dormancy phase, a decrease of total NSCs was recorded, driven by a severe decrease of starch although associated with a simultaneous increase in soluble sugars.

**Figure 3:**
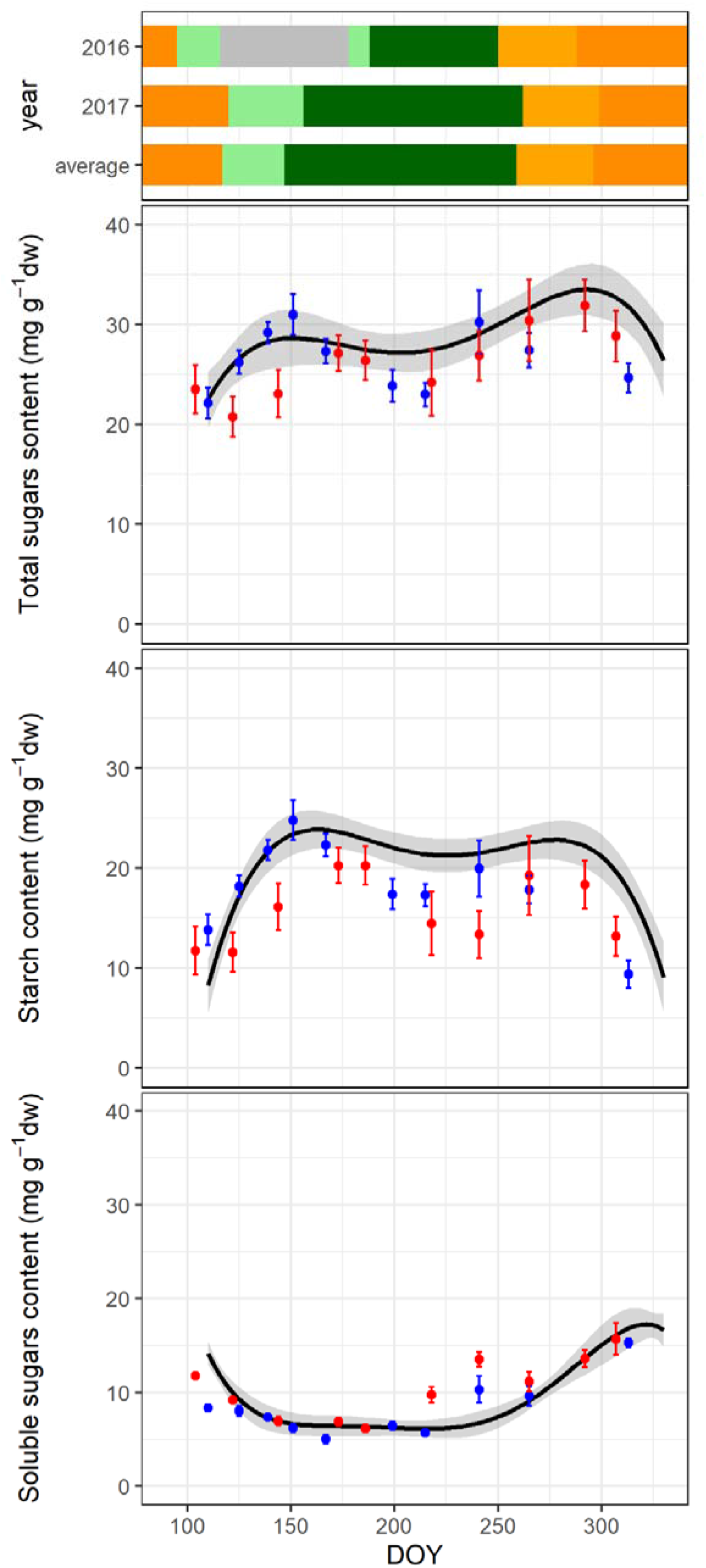
Phenological data for the experimental beech forest site (top panel) and seasonal dynamic of NSCs content as total NSCs (panel *a*), starch (panel *b*) and soluble sugars (panel *c*).In the top panel, the different colours represent dormancy (dark orange), the period between the green up and the maximum Leaf area index (LAI, m^2^ m^−2^) value (light green), the maximum LAI (dark green), the senescence phase (light orange) and, finally, the leafless period after the late frost in 2016 (grey). In the panels a, b, and c, blue and red dots represent carbohydrate concentrations of 2016 and 2017, respectively, while the black lines and grey area show modelled intra annual dynamic of carbohydrates and interval of confidence, respectively. Each point is the mean of five beech trees and bars are the standard errors (see Material and Methods). Modelled values are derived from 39 and 28 measurements of starch and soluble sugar content, respectively.

**Table 1:** Parameters of the polynomial model describing the intra-annual variation of non-structural carbohydrates (total sugars, starch and soluble sugars content) in beech wood, n is the number of samplings used for the analysis and RMSE (mg g _DW_^−1^) is the root mean square error.

| Components | yo | a | b | c | d | e | n | R<sup>2</sup> | RMSE | p-value |
| --- | --- | --- | --- | --- | --- | --- | --- | --- | --- | --- |
| Total sugars | 28.29 | 11.21 | -5.47 | -3.68 | -10.23 | 4.95 | 28 | 0.64 | 2.36 | <0.001 |
| Starch | 19.55 | 4.98 | -26.09 | 2.43 | -14.76 | - | 39 | 0.78 | 2.57 | <0.001 |
| Soluble sugars | 9.41 | 8.13 | 20.28 | -3.31 | 1.72 | -3.97 | 28 | 0.93 | 1.00 | <0.001 |

In 2016, the lowest soluble sugars content (5.02 ± 0.46 mg g _DW_^−1^) was measured close to the build-up of the new photosynthetic apparatus. In the same year, the maximum soluble sugars content (15.29 ± 0.48 mg g_DW_^−1^) was measured at the end of the vegetative season, during the dormancy phase, while two peaks of starch content were measured after the canopy destruction (24.80 ± 0.20 mg g _DW_^−1^) and close to the beginning of the senescence phase (19.93 ± 2.82 mg g _DW_^−1^).

In 2017, the lowest content of soluble sugars (6.16 ± 1.36 mg g _DW_^−1^) was measured at Day of the Year (DOY) 186, while the lowest starch contents were measured before the green up (13.82 ± 1.51 mg g _DW_^−1^) and during the dormancy (9.37 ± 1.36 mg g _DW_^−1^).

Although the seasonal trends of carbohydrates accumulation in wood samples in 2016 and 2017, were similar to the modelled NSCs dynamic recorded in the reference period, some substantial differences can be observed. In 2016, the higher starch content, balanced by a low soluble sugars amount, was recorded soon before the green up. After the second leaf re-sprouting, starch content decreased considerably, reaching a value lower than the modelled reference value at the site. In August 2016, both starch and soluble sugars increased until leaves senescence, which occurred earlier than the average of the site. After that, a reduction of the total carbohydrate reserves was observed.

The lower amount of storage carbohydrates reached in 2016, directly affected the starch and total NSCs amount recorded during the first part of the vegetative season in 2017, when a very low content of starch and total carbohydrates was measured. At the end of July 2017, although the starch content was lower than the modelled value of the site, a refilling of total carbohydrate reserves was observed. The drought stress event of August 2017 strongly affected the composition of carbohydrate reserves due to a severe starch hydrolysis, leading to a decrease of starch content of about 35% and a parallel increase of soluble sugars. During the late phase of vegetative season of 2017, the carbohydrates pattern returned close to the modelled intra annual dynamic, although with a limited reduction compared to the site average.

## Discussion

### The buffering capacity of NSCs in response to the late frost

As already observed at the site and as reported in the literature, the seasonal dynamics of NSCs play a crucial role in regulating C source-sink balance through buffering the difference between C supply and demand (Scartazza *et al.*, 2013; Fatichi *et al.*, 2014; Collalti *et al.*, 2018, 2020). The complete destruction of photosynthetic canopy and the strong reduction of stem radial growth during springtime 2016 (May-June) following the frost event, were associated with an increase of stemwood NSCs due to starch accumulation. An increase in total stemwood NSCs has been previously observed from November to March in other temperate forests and it was attributed to remobilization of sugars from storage compartments in coarse roots in advance of the C demands associated with springtime growth (Hoch *et al.*, 2003; Hartmann & Trumbore, 2016). The NSCs seasonal dynamic shows that starch accumulation in beech occurs during the formation of the new crown, in the presence of the potentially dominating sink represented by new growing leaves and shoots, while soluble sugars are decreasing. Furthermore, our results confirm that the accumulation of starch in sapwood of beech trees during springtime is not necessarily supported by freshly produced photosynthates. In 2016, it occurs, uniquely, as the result of the remobilization of already existing soluble NSCs, including those remobilised from below-ground organs. The normal starch rise in spring could be favoured by the destruction of the developing canopy leaves. This condition leads to a high concentration of soluble sugars within the stemwood that, concurrently to the springtime increased air temperatures, favour synthesis of starch over its degradation (Witt & Sauter, 1994). Indeed, it was recently demonstrated in one-year old shoots of *Juglans regia* L. that wood accumulation of starch can be increased when photosynthate export from the shoot is blocked by girdling. In such case, the increase of starch accumulation is accompanied by an increase of the total activity of ADPglucose pyrophosphorylase (Moscatello *et al.*, 2017), an enzyme well known for its high control over starch synthesis. Thus, the spring programmed activation of starch synthesis in wood can occur even when C resources are very limited by the absence of a photosynthesizing crown. This strongly supports the hypothesis of an active control of the accumulation and buffering role of NSCs in wood (Sala *et al.*, 2012; Collalti *et al.*, 2020).

The buffering role of NSCs to compensate the difference between C sink and C supply was also particularly evident during the late spring and early summer 2016, when stemwood starch reserves were partially hydrolysed for sustaining the second leaf re-sprouting, causing a slight decrease of NSCs during July compared to the modelled values of the site. D’Andrea *et al.* (2019), using the ^14^C bomb technique, found that up to 5 years old reserves can be mobilized to sustain the tree metabolic activities and leaf re-sprouting after the frost damage, further supporting the role of NSCs in the resilience of beech to extreme weather events such as late spring frost. This information is crucial in understanding the resilience capacity of Southern beech forest to late frost, especially considering that extreme warm events may have particularly strong influences at the end of winter when some species interrupt dormancy and the risk of freezing remains relatively high (Ladwig *et al.*, 2019). After the complete canopy defoliation subsequent to the frost event, cambium activity was maintained at low rates, representing an additional C sink during the leafless period fuelled by the stemwood reserves (D’Andrea *et al.*, 2020b). During the second part of the season (August-September), the new assimilates from the canopy are mainly used to sustain C sink activities related to wall thickening and lignification phase (Prislan *et al.*, 2018) and to refill the starch reserves within the stemwood. However, C allocation to cell wall thickening, was extremely limited in 2016 due to strong reduction of xylem cells production (D’Andrea *et al.*, 2020b). This possibly led to the increase of both starch and soluble sugars observed in stemwood of beech plants in August 2016, notwithstanding the reduction of the net ecosystem exchange (NEE) compared to the 2000-2015 values (Bascietto *et al.*, 2018). The NEE decrease in 2016 is related to canopy destruction after the late frost and, overall, to the lower LAI, likely associated also with a reduction of photosynthetic efficiency of defoliated beech trees (Gottardini *et al.*, 2020). In that year, the strong reduction of sink activity, concomitantly with the seasonal decrease of air temperatures, could contribute to the slightly anticipated closure of the season. After leaf shedding, starch was partly hydrolysed and converted to soluble sugars to reduce cell osmotic potential and induce cold tolerance (Bonhomme *et al.*, 2005; Tixier & Sperling, 2015).

### The role of NSCs to face the summer drought

The slight reduction of C reserves at the end of the 2016 growing season impacted the dynamic of the following year. Notwithstanding that the content of starch showed the typical seasonal trend of the site, the starch content in woody tissue from bud break till the end of June 2017 was clearly lower than the modelled reference NSCs dynamic of the site, while there were not relevant differences among modelled and measured content of soluble NSCs. This phase was followed by a sharp rise in the stem radial growth as typical for the site (Scartazza *et al.*, 2013). In summer 2017, the warm drought event had a strong effect on NSCs dynamic, leading to starch hydrolysis and accumulation of soluble sugars in woody tissue. As drought induces a partial stomatal closure that reduces C uptake, trees are forced to depend more on NSCs storage to sustain metabolic activities, defence mechanisms against pathogens and osmoregulation processes (McDowell, 2011; Hartmann & Trumbore, 2016). The negative effects of drought can be exacerbated by the concomitant temperatures higher than average (i.e. the so-called “hot” drought), strongly affecting water transport between roots and canopy (Hartmann & Trumbore, 2016). Moreover, respiration increases with temperature leading to NSCs depletion (Guidolotti *et al.*, 2013; Collalti *et al.*, 2018), while, at the opposite, drought alone reduces whole-plant and root respiration (Hartmann *et al.*, 2013). The observed increase of wood soluble sugars concentration during July-August 2017 is in agreement with the key role of these C compounds as compatible solutes for osmoregulation (Chaves *et al.*, 2003). Indeed, plants under drought conditions can actively control the osmotic cell pressure to avoid tissue dehydration and maintain the physiological functions by increasing the concentration of different kinds of compatible solutes such as betaines, amino acids and sugars (Morgan, 1984). In our study, the increased concentration of stemwood soluble sugars during drought was due to both hexoses (glucose and fructose) and sucrose (data not shown), according to previous findings (Fu & Fry, 2010; Yang, 2013). In addition, NSCs have also a relevant role to maintain xylem transport and embolism repair under drought conditions (Scartazza *et al.*, 2015; Hartmann & Trumbore, 2016). The so called ‘*C starvation hypothesis’* (Mcdowell *et al.*, 2008) speculates that the drought-induced stomatal closure minimizes hydraulic failure but, at the same time, causes a decline of photosynthetic uptake, possibly leading to C starvation as carbohydrates demand continues for the maintenance of metabolism and defence. Moreover, the concomitance of elevated temperatures could accelerate the starch depletion leading to tree mortality (Adams *et al.*, 2009), suggesting that trees, to avoid this risk, should be able to maintain a minimum (safety) level of reserve under drought and warm conditions (McDowell & Sevanto, 2010). Our results support this hypothesis, showing that, notwithstanding the partial starch hydrolysis, the total NSCs contents were only slightly affected, indicating that beech trees were able to counteract a relatively brief and intense “hot” drought event by the interconversion between starch and soluble sugars without drastically affecting the total C storage reserves in woody tissue. This extreme weather event delayed the starch accumulation in woody tissue during the late summer-autumn period, when storage carbohydrates represent one of the major forest C sinks (Scartazza *et al.*, 2013). However, at the end of the 2017 vegetative season, trees were able to store similar amounts of starch and total NSCs compared to the modelled reference value of the site, confirming that the studied forest showed an efficient internal regulation mechanism able to respond to environmental factors with short-to medium-term homeostatic equilibrium (Scartazza *et al.*, 2013; Dietrich *et al.*, 2018). The absence of a strong depletion of NSCs even during two sequential years characterised by extreme weather events that strongly reduced C supply and, at the same time, increased C demand for sustaining stress-recovery (frost) and stress-tolerance (drought) processes, further support the hypothesis that C reserves in plants can be tightly actively managed. In this view, wood NCS synthesis, cleavage, interconversion, mobilisation and allocation need to be actively controlled at the physiological biochemical and molecular level, to optimize growth and survival in the long-term (Sala *et al.* 2012; Collalti *et al.* 2018; Collalti *et al.* 2020).

### The interplay between carbon reserves and growth

At the Collelongo-Selva Piana beech forest, after leaves development, stem radial growth represents the main C sink during the first part of the vegetative season reaching the maximum rate in July. Along the season in the late summer and early autumn period the photosynthates are mainly allocated to the storage C compounds in woody tissues, as described by the models of carbohydrates intra-annual dynamic and by previous studies (Michelot *et al.*, 2012; Scartazza *et al.*, 2013).

The complete canopy destruction in spring 2016 led to a drastic reduction of radial growth without affecting stemwood C reserves dynamic during the first part of the vegetative season, supporting the hypothesis that stem radial growth of diffuse-porous tree species starts soon after leaf expansion and C needs are more likely to be supplied by the new assimilates rather than from the C pool stored within the stem sapwood (Barbaroux & Bréda, 2002; Čufar *et al.*, 2008; Zein *et al.*, 2011; Michelot *et al.*, 2012). After the “second” green up in July 2016, the accumulation of C reserves was prioritized over allocating recently fixed C to stem growth.

In 2017, at the beginning of vegetative season, the new assimilates produced by the canopy photosynthesis were mainly used for sustaining the stem radial growth, which, differently from 2016, reached values of BAI similar to those observed for the reference period (1989-2015). In 2017, the extreme summer drought affected NSCs dynamic but had only very limited effects on annual stem radial growth, as already observed for other tree species growing in the Mediterranean area, which adopt a stress avoidance strategy, adjusting the end of xylem growth before potential stressful conditions may occur (e.g. Lempereur *et al.* 2015; Forner, Valladares, Bonal, Granier & Grossiord 2018). Ultimately, our results confirm that for Mediterranean beech, growth is more negatively impacted by spring late frost than summer drought (Gazol *et al.*, 2019).

Moreover, our results suggest that in long-term adapted mature beech forests, summer drought has limited effects on stem growth, being it mainly dependent on new photosynthates produced in spring. At the opposite summer drought was reported to have important effects on stem growth of young beech trees (Chuste *et al.*, 2020).

In beech trees, the duration of wood formation was found to be positively correlated with increasing latitude, with warmer and drier conditions reducing the length of xylogenesis (Martinez *et al.*, 2016). We speculate that the shorter period of wood formation in the more drought-prone Mediterranean forests, could be also related to a local adaptation of beech to environmental constraints (Jump & Peñuelas, 2007). The reduced sink activity (related to radial growth, wall thickening and lignification) during extreme weather events could be functional to prevent NSCs depletion (Anderegg, 2012; Dietrich *et al.*, 2018). The maintenance of high NSCs concentration and control over NSCs metabolism (e.g. starch hydrolysis) during severe drought events contribute to avoid xylem hydraulic failure and strong damages, as observed conversely in Central European beech (Schuldt *et al.*, 2020). It should be noted that NSCs, including starch, can be rapidly interconverted, ensuring a rapid hexose supply to the hexose phosphate pool. The hexose phosphate pool then supports both metabolic and structural cell requirements for reduced carbon, ranging from glycolysis and respiratory metabolism to cell wall polymer synthesis. On the contrary, assimilates ending up in cell wall components cannot be reclaimed for metabolism, representing, in this respect, almost a dead end, at least in the short period. Hence, under photosynthate famine conditions, prioritization of photosynthate allocation to NSCs, might ensure the maintenance of a sufficient amount of metabolically available reduced carbon. This acclimatory choice seems more conservative than supporting end point-like allocation of photosynthates to cell wall components and ensure a much higher plasticity to support plant response to environmental constraints. Furthermore, allocation of photosynthates to NSCs is less energy costly than building new cell walls polymers (Rodríguez-Calcerrada *et al.*, 2019) again making this choice more conservative in case of reduced source activity. The ability of Mediterranean beech trees to store C reserves also during stressful years could represent an active strategy for optimizing growth and survival and coping with the increasing frequency of extreme weather events.

Summarizing, our study elucidated the mechanisms connected to the impact of late frost and summer drought on sink processes (stem and foliage growth, allocation to reserve pool) in a Mediterranean beech forest. Synthesis, cleavage, interconversion, mobilisation and allocation of wood NSCs are all finely regulated processes and play a key role in counteracting the negative effects of both late frost and summer drought, ensuring plant survival and buffering the difference between C supply and demand under extreme weather event conditions. This information suggest that C reserves could be crucial for resilience of beech because of the expected increasing frequency of extreme weather events under the future global changes and may be useful for adaptive future management strategies of beech forests in the Mediterranean area and Europe.

## Acknowledgements

The activities of Negar Rezaie at the wood anatomy laboratory of Slovenian Forestry Institute were supported by an Excellence Research Award of the National Research Council of Italy, Department of Biology, Agriculture, and Food Secures (Prot. 71951, 06/11/2017). Collelongo-Selva Piana is one of the sites of the Italian Long-Term Ecological Research network (LTER-Italy), part of the International LTER network (ILTER). Research at the site in the years of this study was funded by the eLTER H2020 project (grant agreement no. 654359).

## Author contribution

E. D’A., A.S, S.M., N.R., G.M. contributed to the design of the research. Fieldwork was carried out by E. D’A., N.R.. Soluble Sugars Content analysis were performed by S.M., A.B, S.P, and A.S.. Data analysis was done by E. D’A, A.S., S.M., data interpretation by all co-authors. The manuscript was written by E. D’A, A.S., S.M., A.C.. All authors read and commented the manuscript.

## Supporting Information

**Table S1**: Dataset of soluble sugar (glucose, fructose and sucrose), starch and total non-structural carbohydrates.

